# SanPy: A whole-cell electrophysiology analysis pipeline

**DOI:** 10.1101/2023.05.06.539660

**Authors:** Laura Guarina, Johnson Tran Le, Theanne N Griffith, Luis Fernando Santana, Robert H Cudmore

## Abstract

The analysis of action potentials and other membrane voltage fluctuations provide a powerful approach for interrogating the function of excitable cells. Yet, a major bottleneck in the interpretation of this critical data is the lack of intuitive, agreed upon software tools for its analysis. Here, we present SanPy, a Python-based open-source and freely available software pipeline for the analysis and exploration of whole-cell current-clamp recordings. SanPy provides a robust computational engine with an application programming interface. Using this, we have developed a cross-platform graphical user interface that does not require programming. SanPy is designed to extract common parameters from action potentials including threshold time and voltage, peak, half-width, and interval statistics. In addition, several cardiac parameters are measured including the early diastolic duration and rate. SanPy is built to be fully extensible by providing a plugin architecture for the addition of new file loaders, analysis, and visualizations. A key feature of SanPy is its focus on quality control and data exploration. In the desktop interface, all plots of the data and analysis are linked allowing simultaneous data visualization from different dimensions with the goal of obtaining ground truth analysis. We provide documentation for all aspects of SanPy including several use cases and examples. To test SanPy, we have performed analysis on current-clamp recordings from heart and brain cells. Taken together, SanPy is a powerful tool for whole-cell current-clamp analysis and lays the foundation for future extension by the scientific community.

**News and Noteworthy:** Whole-cell current-clamp recording is a critical technique to understand detailed biophysical mechanisms at the cellular and network levels. Yet, the analysis of this data is by no means standardized. Here, we present SanPy, a Python based open-source software pipeline for the exploration and analysis of current-clamp recordings. SanPy provides a powerful computational engine with an application programming interface. Finally, SanPy provides a cross-platform point-and-click graphical user interface that does not require programming.

## Introduction

Patch-clamp electrophysiological recordings are an invaluable tool to examine detailed biophysical properties and mechanisms at the cellular level with a high signal-to-noise ratio and exquisite temporal resolution (1, 2). In the whole-cell configuration, a recording is performed by placing a small (µm scale) glass pipette on a cell membrane to form a high-resistance giga-Ohm seal, and then electrical access to the inside of the cell is achieved either by rupturing (with pressure) or perforating (with anti-biotics) the membrane under the pipette. Once whole-cell access is achieved, either currents or voltages can be recorded and delivered to the cell.

In whole-cell current-clamp mode, the membrane voltage of a cell is recorded, and a range of hyperpolarizing and depolarizing currents can be delivered to characterize a cell’s response. Whole-cell current-clamp recording, and analysis is not limited to a particular cell type but of general interest are excitable cells such as cardiac myocytes and brain neurons. The common feature of these cell types is they possess regenerative potentials termed action potentials (APs). APs are mediated by both the kinetics and voltage dependance of currents produced by the opening and closing of ion channel membrane proteins including but not limited to Na^+^, K^+^, Ca^2+^, and Cl^-^ channels.

The AP pattern, frequency, and kinetics collectively define what is referred to as the intrinsic excitability of a cell (3, 4). Using current-clamp to examine intrinsic excitability allows a detailed understanding of cellular and network function. For example, in cardiac myocytes, the AP frequency and its reliability can be used to predict heart rate and the potential for arrhythmias. In neurons, the frequency of APs in response to a given current injection can be used to predict how a neuron will integrate synaptic input to generate AP output. This has important implication for understanding, for example, sensory encoding and information transmission in networks of neurons.

Using whole-cell current-clamp electrophysiology to understand intrinsic excitability has wide ranging implications in the development of detailed biophysical models of cell function, plasticity, and disease. In neurons, the plasticity of intrinsic excitability has been examined using both *ex vivo* and *in vivo* stimulation (5–7). It has also been shown that *in vivo* sensory experience and chronic activity regimes can sculpt neuronal excitability (8–12). Examining intrinsic excitability is also critical in understanding functional differences between different classes of neurons (13, 14). In the heart, the intrinsic excitability of cardiac myocytes has been examined with a complimentary set of hypotheses. For example, it has been used to examine the differences in the excitability between anatomically distinct regions of the sinoatrial node and how heart failure can sculpt cardiac myocyte excitability (15, 16).

The majority of current-clamp analysis is performed using either commercial software such as pClamp (Molecular Devices) or SutterPatch (Sutter Instruments) or implemented as scripts in general purpose software such as Matlab (Mathworks) or Igor Pro (Sutter Instruments). Because of the inherent heterogeneity in these analysis methods, it is often difficult to compare the results between studies. A major bottleneck to comparing different analysis is there is still no agreed-upon standard. Such a standard would better allow for the comparison of results between different groups, individuals within a group, and different preparations.

With the advent of open-source programming languages such as Python, we now have the tools in hand to ensure the development of community-based analysis software (17, 18). The National Institute of Health (NIH) has been progressively implementing requirements that NIH funded projects make all raw data and analysis publicly available. This was most recently codified with the 2023 NIH Data Management & Sharing Policy. The development of analysis software in an open-source language such as Python has several benefits including the ease of implementation and extensibility with scientific computing packages such as NumPy, Pandas, and SciPy (19–21). A few current-clamp analysis packages have been developed in Python including paramAP (22) and StimFit (23) but to the best of our knowledge, these are scripting packages with no graphical user interface (GUI), thus limiting their adoption by the broader biological research community.

Here, we present SanPy, an extensible and fully documented open-source pipeline for electrophysiology analysis written in Python. SanPy includes an easy-to-use desktop GUI that requires no programming coupled with a backend application programming interface (API) for scripting. SanPy will calculate common AP metrics such as the time and voltage threshold, peak, half-width, and interval statistics. In addition, SanPy can analyze these AP parameters in response to varying levels of injected current. Because SanPy provides a GUI, it can be used to interactively interrogate and adjust the detection parameters and resulting analysis to arrive at the ground-truth. The SanPy API can easily be extended with plugins to include loading from any raw data file format, new analysis, and novel visualizations tailored to a particular experiment’s needs. To test SanPy on real world data, we provide example analysis in both cardiac myocytes, and central- and peripheral nervous system neurons. To facilitate the adoption of SanPy, we have included several examples that show how to add to its core features.

## Methods

All code is written Python, is open-source, and available on GitHub (https://github.com/cudmore/sanpy) (24). One file downloads are provided for the desktop GUI application that runs on both macOS and Windows. These click-and-run desktop GUI applications do not require any programming experience or additional system installation. For command line installation, a Python package, sanpy-ephys, is available on PyPi (https://pypi.org/project/sanpy-ephys/). All documentation can be found in the online SanPy documentation (https://cudmore.github.io/SanPy/).

### Release Engineering

To ensure reproducibility, each time a new version of SanPy is released it includes a unique version number. This version number is saved in all output files. Because SanPy is archived on GitHub, any previously released version can be run to compare any differences in the results. SanPy has been developed using several best engineering practices (25) including continuous integration using GitHub Workflows to run tests (pytest) on the code to ensure it generates pre-defined and expected results and to verify the code functions on a matrix of end-user machines including macOS, Windows, and Linux as well as for a number of Python versions starting with Python 3.8 and currently extending to Python 3.11.

### Code design

All code has been designed to be modular by separating the computational API from the frontend GUI. The API is a Python class library that provides functionality to load raw data, perform analysis, and save the results. This API allows programmatic control of all SanPy functionality without requiring interface libraries. This allows SanPy to interoperate with other software packages and to run headless either on a local machine or in the cloud. The API uses a number of standard Python packages including (but are not limited to): NumPy, Pandas, SciPy, pyAbf, and h5py (19–21, 26, 27).

### Desktop GUI

The desktop GUI is cross platform and will run on macOS, Microsoft Windows, and Linux operating systems. The GUI is implemented in Python using packages such as PyQt, PQtGraph, Matplotlib, and seaborn (28, 29). The desktop GUI is built and distributed as a macOS app and a Microsoft Windows exe using the Python package PyInstaller.

The GUI is split into independent widgets such as file list, raw data plot, and plugins like a scatter plot and tabular results. Each widget communicates the state as a user interacts and passes messages to other widgets. This system is referred to as a publisher-subscriber architecture. With this, selections in one widget will be propagated to all other widgets, making the user interface highly interactive and exploratory. Because of this architecture, individual widgets need not be aware of other widgets but instead emit signals on state changes so other widgets can connect to receive state changes to respond themselves.

### Loading raw data and saving analysis

Raw data can be loaded from a number of raw data formats. To ensure SanPy can be used with raw data from a wide-range of acquisition system and file-formats, we have implemented a file-loader plugin architecture allowing users to implement custom functions to load any file format (**See Supplemental Recipe 1**). Using this architecture, SanPy includes file loader plugins for Axon binary (abf) and text (atf) files (using the pyAbf Python package), comma-separated-value (CSV) text files, and Matlab (Mathworks) files. Internally, all analysis is saved in an HDF5 file format using the Python package h5py. Example code is provided to load and perform additional analysis on the natively saved HDF5 files. Finally, all analysis can be saved to a CSV for additional analysis in any programming language or desktop application.

### Extending the analysis of SanPy

A plugin architecture is provided such that users can write their own analysis code to extend the core analysis provided by SanPy. This custom analysis is automatically incorporated into the GUI and is saved and loaded with no additional customization. Several examples are provided to show how the analysis can be extended (**See Supplemental Recipe 2**).

### Extending the SanPy GUI

A plugin architecture is provided that allows users to extend the GUI to meet their particular analysis needs. This plugin architecture uses class inheritance from a base plugin class. With this inheritance, the user is given access to all the raw data and analysis displayed in the core GUI as well as signals which allow plugins to dynamically interact with the state of the SanPy GUI (**See Supplemental Recipe 2)**. We provide a number of plugins that are built-in to the core of SanPy (**Table 1**).

**Table 1.** Provided plugins. Two main categories of plugins are provided including (i) plot and (ii) table.

| Plugin Name | Type | Description |
| --- | --- | --- |
| Plot Recording | Plot | Plot the raw recording with all analysis results overlaid |
| Spike Clips | Plot | Overlay APs aligned to their threshold. Includes phase plots and waterfall plots' |
| Plot Scatter | Plot | Plot scatter plots to examine the relationship between analysis results. |
| Plot FI | Plot | Plot summary analysis for a typical experiment delivering a range of current injection amplitudes. |
| Plot Analysis | Plot | Examine multiple plots at the same time to explore the relationships between parameters. |
| Export Trace | Plot | Visually edit the zoom and labelling of a raw trace to export to a file for publication. |
| FFT | Plot | Performs both FFT and PSD analysis on raw data. This includes filtering signals with Bessel filtering. |
| Detection Parameters | Table | Set all detection parameters, load presets, and create user defined subsets. |
| Summary Analysis | Table | a) Show a tabular summary of all detected spikes and all analysis results.<br>b) Show a tabular summary of analysis results with the mean, std, sem, and n for each measurement.<br>c) Show a tabular summary of all detection errors. |
| SanPy Log | Table | A tabular summary of the SanPy log. This is a "lab notebook" of actions taken on the SanPy interface. This can be sent to developers to troubleshoot and debug issues. |

### Signal detection algorithm and analysis results

To capture the wide range of cell types with potentially different AP phenotypes and membrane time-constants, SanPy has a flexible set of detection parameters (**See Supplemental Table 1**). A number of pre-set detection parameter sets is included for common cell types such as ventricular and sinoatrial node cardiac myocytes as well as fast- and slow-spiking neurons. We also provided detection parameters for sub-threshold events such as synaptic currents and Ca^2+^ sparks.

Optional preprocessing of raw data is performed with either Median or Savitzky-Golay filters. These were chosen as they preserve the timing of transients such as the temporal onset and shape of an AP.

Initial AP detection is done with a threshold in one of two ways, either a threshold in the first derivative of the membrane potential (dV/dt) or a voltage threshold in the primary recording (mV). The dV/dt threshold is the desired detection strategy as it results in a more precise measure of the time of the AP threshold potential (mV).

False positives are removed by examining both the AP waveform and timing with respect to the previous AP. APs are only accepted if they occur on a positive membrane potential trajectory. Rapidly occurring APs are removed with a user specified refractory period (ms). This is useful to remove erroneous APs that occur when, for example, an AP has a very long duration and is potentially noisy. Once the time of each AP is calculated, several parameters are extracted by analyzing the precise waveform within an AP (**See Supplemental Table 2**). This includes, for example, AP peak (mV) and amplitude, pre-AP rise and post-AP decay times (ms), and half-widths (ms). Interval statistics are also extracted such as AP frequency (Hz) and its inverse the AP inter-spike-interval (ms). Several measurements taken from the cardiac myocyte literature are included such as early diastolic duration (ms) and rate (dV/dt) as well as the maximum depolarizing potential between APs (16).

During AP parameter extraction, the algorithms will occasionally fail. This can occur if the recording is relatively noisy or if the detection parameters are mismatched with the actual time-constants of the cell being analyzed. Care has been taken to capture these detection errors and ensure the resulting analysis values do not return erroneous results. All errors are logged and saved with the analysis, allowing users to browse potentially problematic APs and to remove them from the analysis accordingly.

### Model neuron

To test the SanPy analysis algorithms, we implemented a stochastic Hodgkin-Huxley model neuron using the ModelDB accession number 144499 (30). Briefly, the model contains Na^+^ (g_Na_ = 120 mS/cm^2^, E_Na_ = 120 mV), K^+^ (g_K_ = 36 mS/cm^2^, E_k_ = -12 mV), and K^+^ leak currents (g_K,Leak_ = 0.3 mS/cm^2^, E _K,Leak_ = 10.6 mV) with a cell capacitance (1 µF/cm^2^). Stochasticity was implemented by adding normally distributed random noise to the subunit variables (*m, h, n*) of the Na^+^ current (31).

### Animals

Male wildtype C57BL/6J mice (The Jackson Laboratory) between 6 and 14 postnatal weeks were used in this study. All mice were maintained, and experiments conducted, in accordance with the University of California, Davis Institutional Animal Care and Use Committee guidelines. Mice were euthanized with an intraperitoneally administered lethal dose of sodium pentobarbital (250 mg/kg).

### Myocyte Dissociation

As described in Grainger et al (2021)(15), following euthanasia, sinoatrial node (SAN) tissue was dissected and placed in Tyrode III solution, containing (in mm) 140 NaCl, 5.4 KCl, 1 MgCl_2_, 5 HEPES, 1.8 CaCl_2_, 5.5 glucose (pH 7.4 with NaOH). The SAN was pinned flat and tissue pieces from the superior and inferior SAN regions (approx. 2–3 mm^2^ each) were harvested and bathed for 5 min at 36°C in Tyrode low Ca^2+^ solution containing (in mm): 140 NaCl, 5.4 KCl, 0.5 MgCl_2_, 0.2 CaCl_2_, 5.0 HEPES, 5.5 d-glucose, 1.2 KH_2_PO_4_, 50 Taurine (pH = 6.9 with NaOH). The tissue sections were then enzymatically digested for 30 min in Tyrode low Ca^2+^ solution (pH 6.9) containing: 9.43 U elastase, 0.89 U protease, 0.27 U collagenase B and bovine serum albumin (1 mg/mL). All solutions used in the cell isolation were maintained at 36°C. Following digestion, tissue segments were rinsed twice with Tyrode low Ca^2+^ solution (pH 6.9) and twice with a Kraft-Brühe solution (4°C ; 80 mml-glutamic acid, 25 mm KCl, 3 mm MgCl_2_, 10 mm KH_2_PO_4_, 20 mm Taurine, 10 mm HEPES, 0.5 mm ethylene glycol-bis(2-aminoethylether)-*N,N,N′,N′*-tetraacetic acid, 10 mm glucose, pH 7.4 with KOH). After 2–3 h at 4°C, the tissue was warmed to 37°C and mechanically dissociated using a fire polished glass pipette.

### Perforated Patch Clamp Recordings

A drop of myocyte cell suspension was placed on a temperature-controlled recording chamber (35–36°C) and cells were allowed to settle and adhere to the chamber for approximately 10 min. Next, the external Ca^2+^ concentration was slowly risen by adding increasing amounts of Tyrode solution to allow a graded transition to physiological Ca^2+^ levels (ie, 1.8 mm). After completion of Ca^2+^ reintroduction, cells were constantly perfused with Tyrode solution.

The internal solution used for current-clamp recordings contained (in mm): 125 K-Aspartate, 10 NaCl, 15 KCl, 1 CaCl_2_, 10 HEPES (pH to 7.2 with KOH). Amphotericin B (100 μm) was added to the internal pipette solution. Fire-polished glass pipettes were dipped in an amphotericin B-free internal solution and then back filled with the amphotericin B–containing internal solution.

Recorded were performed using a patch-clamp amplifier (MultiClamp 700B, Molecular Devices, San Jose, CA) and pCLAMP 10 software (Molecular Devices). Cells were initially patched in voltage-clamp mode and after the formation of a giga-ohm seal required 1–5 min until amphotericin B permeabilized the membrane, allowing electrical access to the cell. The amplifier was then switched to current-clamp mode to record membrane potential (mV). All recordings were acquired at 10 kHz.

Spontaneous APs were recorded during a control period (1-2 min). The beta adrenergic receptor agonist isoproterenol (ISO; 100 nM) was then bath applied. Recordings continued and spontaneous APs were recorded in the presence of ISO.

### Statistics

All data is presented as mean ± standard deviation unless otherwise noted. All statistical tests were performed with the non-parametric Mann-Whitney U test using the Python package SciPy (20). A *P*-value of less than 0.05 was considered significant and all *P*-values are reported as their actual values unless *P* < 0.0005.

## Results

### SanPy desktop application

To use the SanPy desktop application (macOS or Windows), a single file is downloaded and run with a double-click. The desktop application is simple to use and by design parallels the suggested workflow (**Figure 1**). First, a folder of raw data files is loaded and displayed in a table, one file per row (**Figure 1a**). Selecting a file propagates this selection to all open windows and plots. After visual inspection of the raw data, detection parameters are set, and APs are detected (**Figure 1b**). Once detected, analysis results are overlaid on the raw data (**Figure 1c**). Users can zoom into raw data, select sub-sets of APs and this selection is propagated to all other views. Once analyzed, plugins can be used to further explore the raw data and analysis results. This interface is intended to provide a feedback loop between raw data, detection parameters, and the analysis results. Multiple iterations may be required to obtain a set of detection parameters that successfully captures the desired analyses results. The end goal of this iterative interface is to arrive at the ground truth analysis easily and quickly.

**Figure 1.**
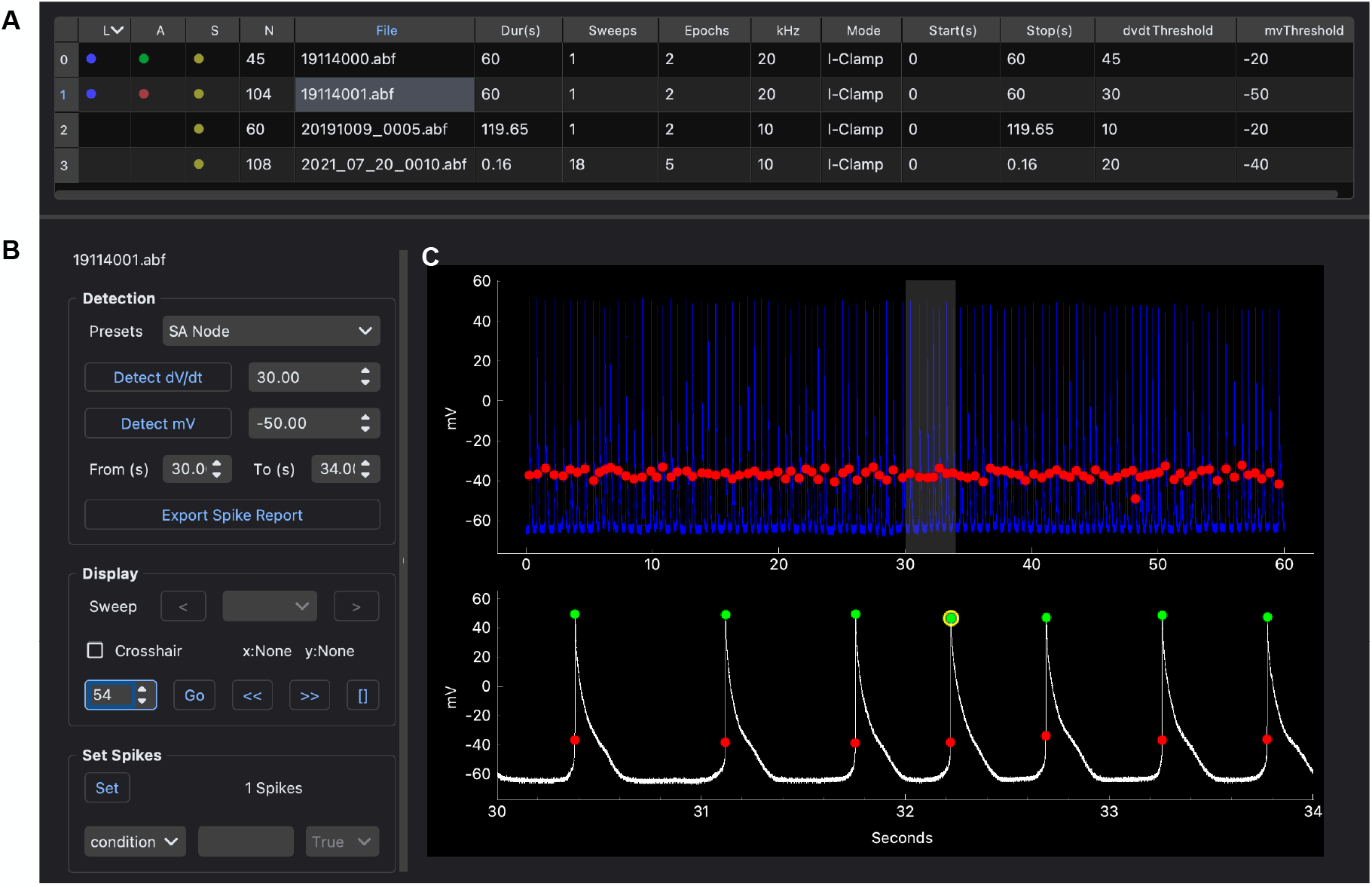
Desktop GUI and workflow. **(A)** A folder is loaded and all raw data files are shown in a table, one file per row. Metadata from each file is shown including the recording duration, the number of sweeps, the sampling interval and the recording mode. In addition, there are columns indicating loaded and analyzed data files including: raw data loaded (column L), analyzed (column A), saved (column S), the number of APs detected (column N), and the number of detection errors (column E). **(B)** Controls to detect and then explore individual APs and analysis results. This includes setting parameters for selected spikes such as their user type and condition. **(C)** Viewing raw data with multiple views. Top plot is the entire raw recording with a selected time-range (gray box). Bottom plot is the zoomed in raw recording. All plots have analysis results overlaid including AP threshold (red circles) and AP peak (green circles). A selected spike is shown in the bottom plot (yellow circle).

SanPy includes per file meta-data to encode a range of features about a recording. This includes information about the recording such as acquisition date and time. There is meta-data for the preparation including species, cell type, postnatal age, sex, and genotype. Additional meta-data is provided to encode experimental conditions. All meta-data is saved with the analysis and is available to all plotting and tabular displays.

All detection parameters and analysis results for each raw data file in a folder or nested folders is saved to a single file (HDF5 format) and is automatically loaded the next time the folder (and its nested folders) is loaded. We chose to save all analysis to a single HDF5 file because these files behave as a random-access database allowing individual pieces, like the analysis for one raw file among hundreds, to be individually saved and loaded regardless of the total number of recordings. Finally, all metadata, detection parameters and analysis results can be exported to several formats including CSV for easy analysis in other software.

### SanPy Plugins

To ensure SanPy can be easily extended, we have created a system where the core functionality can be expanded using a plugin architecture (**See Supplemental Recipe 3)**. A key feature of this system is that it provides a concise API to connect to the raw data and analysis results. The plugin architecture is designed to be down-stream of the core SanPy functionality. With this, any errors in a plugin will not disrupt the functioning of the core SanPy functions. The range of plugins is expanding and there are currently two broad categories including graphical and tabular display of data and its analysis (**Table 1**). Several graphical plugins are shown in **Figure 2**. SanPy is built with a plugin to browse the raw data with all detection parameters overlaid (Plot Recording; **Figure 2a**). There is a scatter-plot plugin to visualize correlations between AP analysis results (Plot Scatter; **Figure 2b**). Finally, a plugin is provided to plot aligned AP clips and to calculate the mean shape of either membrane potential (Vm) or its first derivative in a phase plot (dV/dt) (Plot Clips; **Figure 2c**). Because all plugins are linked back to the main interface (**Figure 1**), SanPy provides a rich GUI to simultaneously explore the raw data and analysis results.

**Figure 2.**
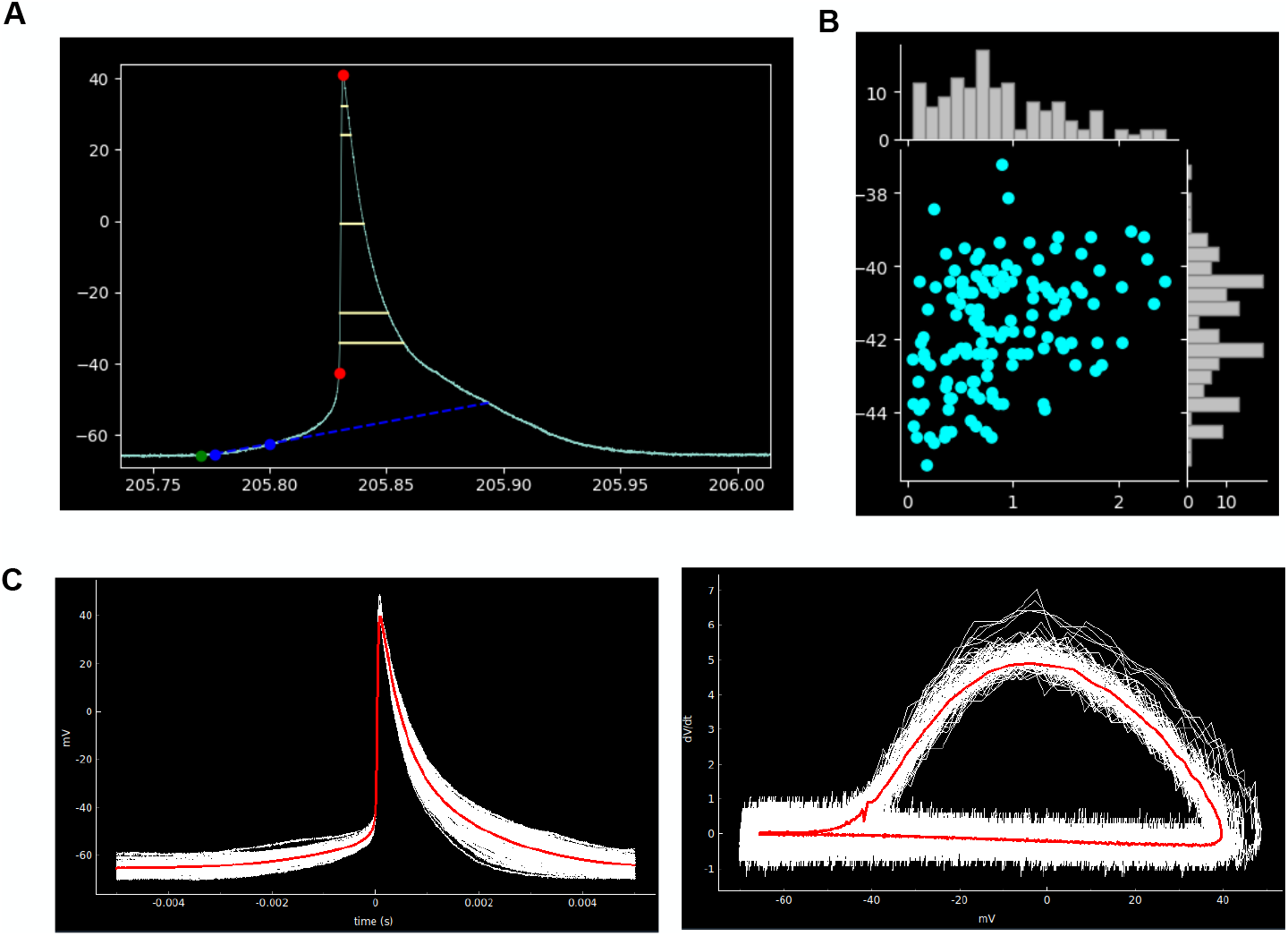
Plotting plugins. **(A)** Plugin to display APs with the analysis results overlaid. Shown here is AP threshold and peak (red circles), maximum depolarizing potential (green circle), early diastolic duration (blue circles), and early diastolic duration rate (blue dotted line), and several AP half-widths representing percent rise of 10,20,50,80,90 % (yellow lines). **(B)** Example scatter-plot plugin showing AP frequency (Hz) versus threshold (mV). The value for each AP is plotted as a symbol (cyan circles). Marginal histograms are showing the distribution of the plotted analysis parameters for both the x and y axes. **(C)** Example of the spike clip plugin showing aligned APs (left panel) and AP phase-plots (right panel). Individual APs are plotted in white and the mean AP waveform across all plotted APs is shown in red.

A canonical current-clamp experiment is to inject a family of current steps with increasing amplitudes and record the AP response to each step. This is commonly used to probe the input-output relationship or transfer-function of a cell. SanPy has several built-in plugins to facilitate the analyze of this class of experiment. Using a recording from a dorsal root ganglion peripheral sensory neuron, we analyzed and visualized the AP response to a family of depolarizing current steps (Plot spikes plugin, **Figure 3a**). To further explore this type of dataset, SanPy provides a plugin to plot all the analysis results as a function of injected current (Plot FI plugin; **Figure 3b**). This plugin will plot any analysis result on the y-axis versus the current injection amplitude and provides a table of summary statistics for a range of derived measurements including the mean, min, max, and coefficient of variation for each current step. As with all SanPy plugins, these results are easily saved as an image file of the plot or as tabular data.

**Figure 3.**
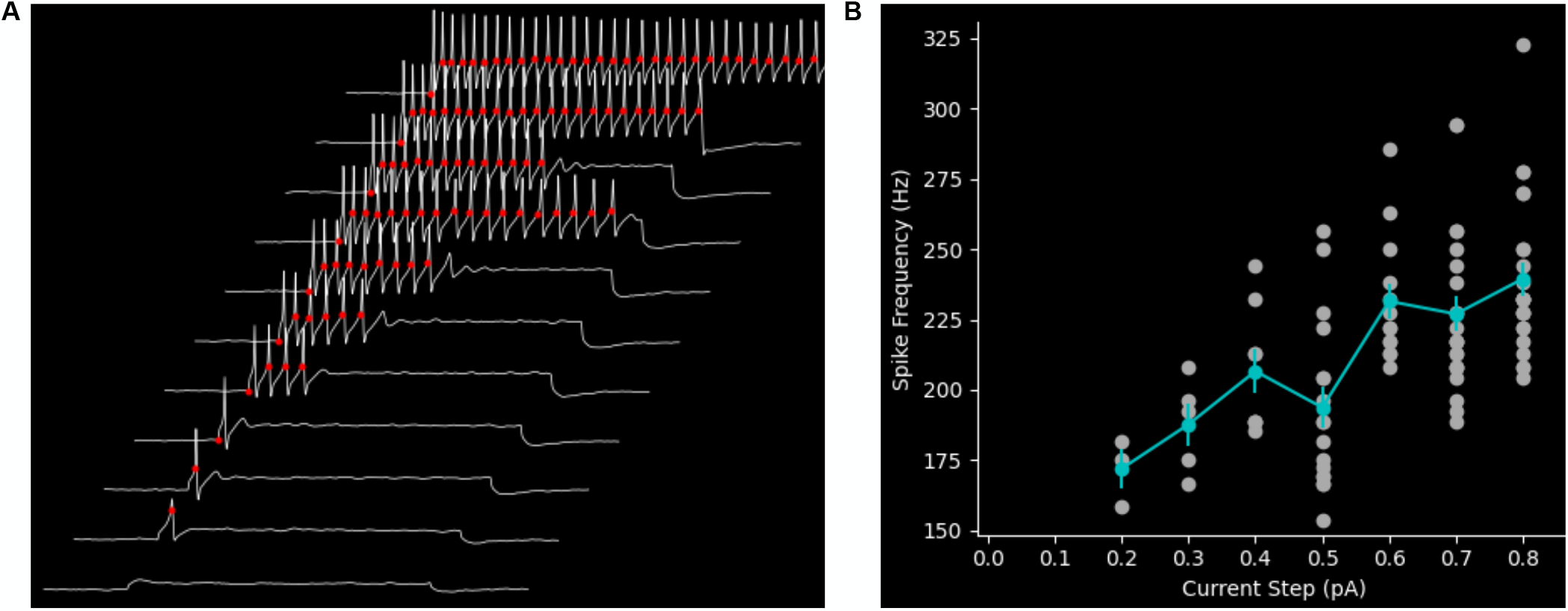
Plugins to analyze the response to a range of current injection amplitudes. **(A)** Plugin to display the response to a family of current injection amplitudes with analysis results overlaid on the raw data. In this example, the response to a family of eleven current injection amplitudes is show. A subset of the analysis results is overlaid on the raw data (AP threshold, red circles). **(B)** Plugin to visualize a summary of the analysis results for an experiment stimulating with a range of current injection amplitudes. Here, as an example, the instantaneous frequency (Hz) for each AP is plotted on the y-axis versus the current injection amplitude (pA). The mean AP frequency for each current step is plotted (cyan markers) with the standard error of the mean (vertical cyan lines).

Once several recordings are analyzed, SanPy provides a powerful pool plot plugin to plot and tabulate analysis across any number of files (**Figure 4**). This plugin effectively allows entire datasets to be visualized and interrogated for trends to test hypothesis. In all plots and tables, data can be grouped by any categorical analysis result (in general, any per file meta-data). Some examples are grouping by sex, genotype, experimental condition, or acquisition date. In addition to scatter (**Figure 4a**) and violin plots (**Figure 4b-c**), other plot styles are provided including line, box, histogram, and cumulative histogram. Finally, each plot has a companion table (**Figure 4c**) that can be copy and pasted into other analysis software.

**Figure 4.**
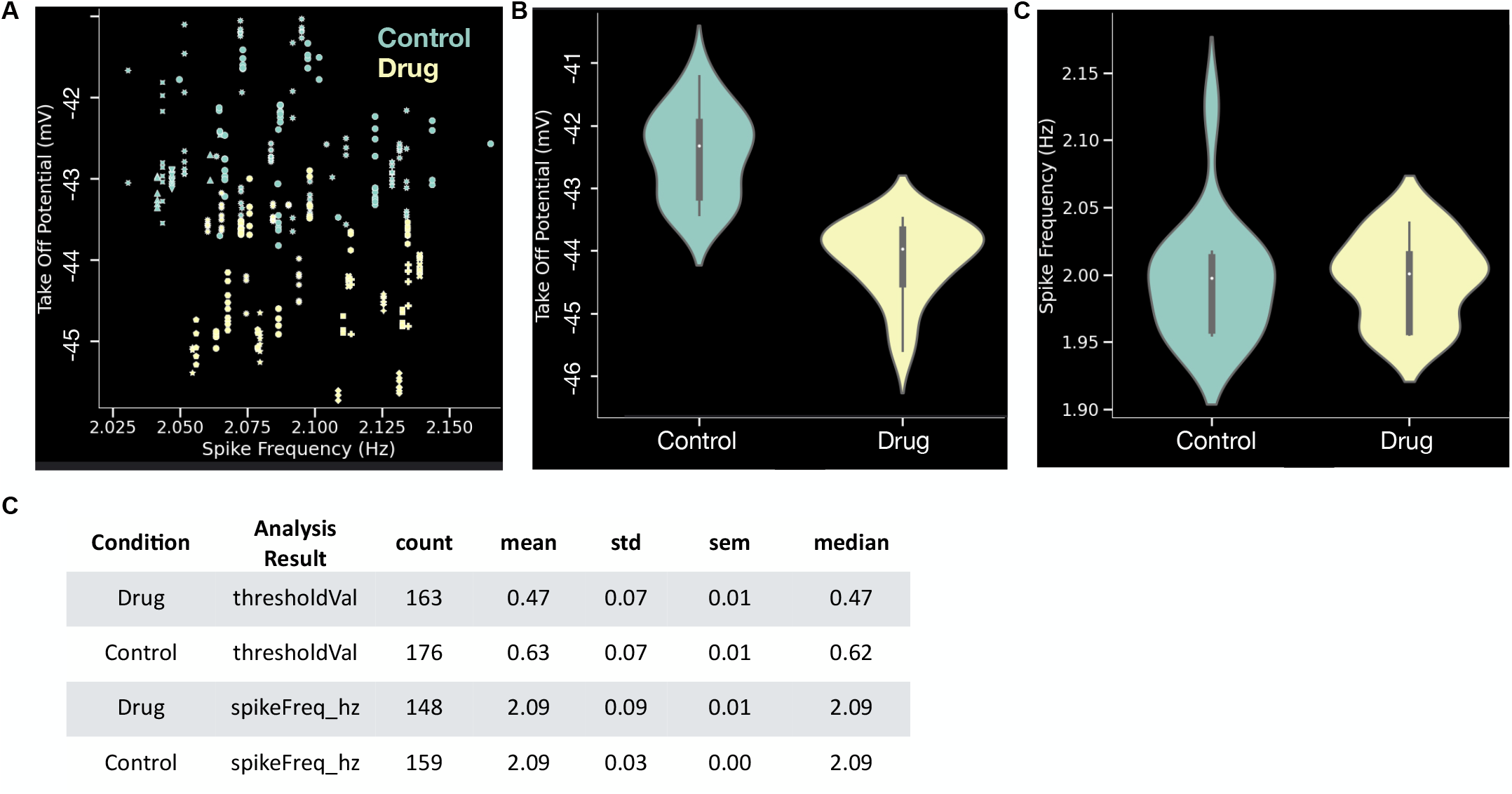
Plugin to pool analysis across files. **(A)** A pooled plot of take off potential (mV) versus spike frequency (Hz). Each symbol is the value for one AP and markers denotes different recordings. Colors denote two different conditions (control and drug). **(B)** Violin plot of take off potential (mV) comparing control to drug conditions. **(C)** Violin plot of spike frequency (Hz) comparing control to drug conditions. (D) Tabular summary of the plots in panels B-C. All plots in these examples are from the same synthetic dataset with 32 recordings and 339 APs.

### Detection Parameters

All detection parameters are rigorously defined by documenting their internal variable name, a human readable name, their units, default values, and a longer description (**Supplemental Table 1**). To capture a range of time-constants in different cell types, we found it necessary to add a number of detection parameters, potentially causing complexity in finding the proper set for high-quality detection within a given recording. To simplify the process for the end-user, we have provided presets for several cell types including ventricular and sinoatrial node cardiac myocytes, slow and fast neuronal cells, and subthreshold peak detection. With the provided detection parameters plugin, user can adjust the predefined detection parameters and once they are satisfactory for a given recording these can be saved and then loaded for future analysis.

### Analysis Results

The most critical function of SanPy is to produce reliable and reproducible analysis results given a raw data recording and a set of detection parameters. To make this as transparent as possible and to allow comparison with other analysis software, we have defined all the SanPy analysis results such that the end-user can understand their meaning and how they were calculated (**Supplemental Table 2**). SanPy generates a range of analysis results taken from both the neuronal and cardiac literature. These include AP time and voltage threshold, peak, and half-width. For cardiac myocyte analysis this includes the early diastolic duration (ms) and rate (mV/s). To ensure SanPy can be used in a wide range of experiments, it can be extended with new analysis results (**Supplemental Recipe 2**).

A key feature of SanPy is that it logs errors during AP detection. For example, when searching for an AP half-width there may be no corresponding falling phase in an AP. In this case, the user must modify the half-width detection parameter. The goal of logging detection errors is to allow the user to fine tune their detection parameters to match their recording. Errors are saved as part of the analysis and each error type is associated with a detection parameter so it can easily be adjusted.

Once APs are detected, SanPy has a set of user configurable notations (e.g., tags) that can be set for each detected AP or for any selection of multiple APs. For example, AP(s) can be tagged as “include” or “exclude”, allowing problematic APs to not be included in the final analysis. This also includes marking an AP or multiple APs with a “user type”. SanPy provides ten different “user type” markings and these can, for example, be used to denote different conditions within a recording such as control and different pharmacological treatments.

### Programming with the API

Users can utilize the SanPy API to work in a familiar environment such as a stand-alone Python script, a Jupyter notebook, or even integrate SanPy functions to interoperate with pre-existing Python packages and applications. A simple example of programming with the API is to write a Python script to load a raw data file, set detection parameters, detect APs, plot the results, and then save all the results (**Figure 5**). This example is using a simulated Hodgkin-Huxley neuron endowed with stochastic Na^+^ channels (see methods). To test the sensitivity of the analysis results, we ran the model with two sets of parameters by modifying the maximal conductance of the K^+^ current (g_K_). This effectively tests a within cell bath application of a drug such as a K^+^ channel blocker. For a 2 second run of the model, for the first half (0-1 s) g_K_ was set to 36 mS/cm^2^ and in the second half (1–2 s) g_K_ was reduced to 30 mS/cm^2^ (a 17% reduction). With this reduction in K^+^ conductance, we hypothesized that AP frequency and half-width would increase while the voltage-threshold for an AP would remain unchanged. As predicted, reducing the maximal conductance of the K^+^ current causes an increase in AP frequency (Control 39.17 ± 17.03 Hz; Drug 55.27 ± 13.48 Hz, p < 0.0005) and half-width (Control 1.14 ± 0.03 ms; Drug 1.22 ± 0.04 ms, p < 0.0005) with no change in the threshold potential (Control - 44.36 ± 0.73 mV; Drug -44.32 ± 0.82 mV, p = 0.91).

**Figure 5.**
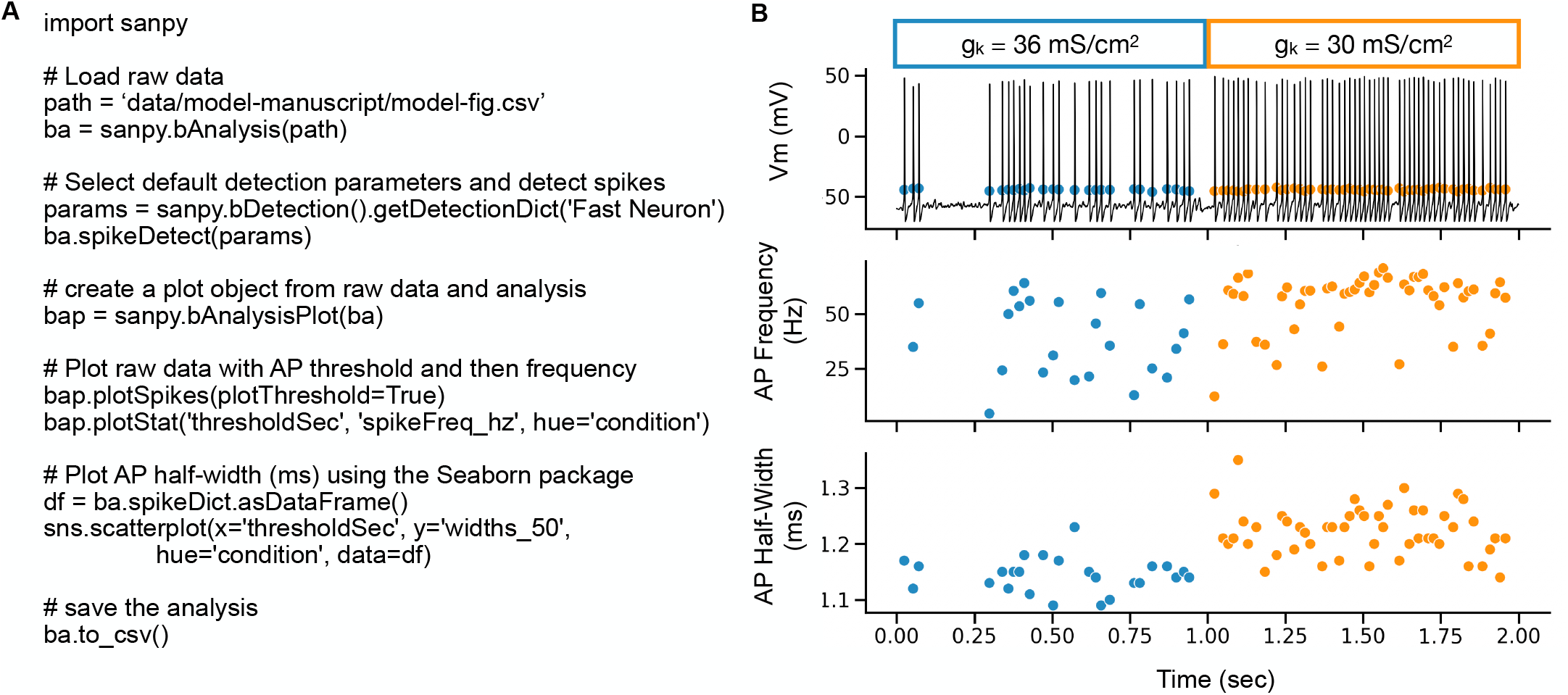
Programming with the API. **(A)** Example Python script to load raw data, detect APs, plot raw data with an overlay of the analysis results, and to save the results to a CSV file. **(B)** Output of the script in (A). Top plot is the raw recording overlaid with AP threshold (mV). Middle plot is AP frequency (Hz) versus time (s). Bottom plot is AP 50 % half-width (ms) versus time (s). Each AP is a symbol (circle). The color of the symbols indicates different conditions within the recording. Here, this represents a change in the maximal conductance of the K^+^ leak channel (g_K_) indicated by blue and orange symbol colors. Please note, all data in this figure is from a stochastic Hodgkin-Huxley neuron model.

### Extending SanPy

A critical component of all analysis software is to provide mechanisms by which it can be extended by a user community. To achieve this, SanPy includes several ways to extend its functionality including API interfaces to (i) create custom file loaders, and (ii) add new analysis to the core detection algorithms, and (iii) create new and novel visualization and tabular plugins.

To ensure SanPy can load raw data from any number of file formats, it includes a plugin architecture to implement user-defined custom file loaders. As an example, SanPy includes custom file loaders for CSV and Matlab files (**Supplemental Recipe 1**).

We are aware that different analysis metrics, measurements, and nomenclatures are used across sub-disciplines such as the cardiac and the neuroscience communities. Thus, additional analysis results can be added by using the provided plugin architecture to encapsulate new analysis and detection measurements. With this system, new measurements from the raw data are seamlessly integrated into the existing SanPy GUI and are automatically saved and exported. As an example, SanPy includes custom analysis code to detect the maximal diastolic depolarization between AP (for cardiac cells) and the 20-80% AP rise time (for both neurons and cardiac cells)(**Supplemental Recipe 2**).

### Benchmarking the SanPy detection algorithm

An important consideration for all analysis software is to determine if the algorithms will scale for high throughput. To examine the time it takes to perform AP detection, we ran AP detection for progressively longer times within a 500 second recording (10 kHz) from an acutely isolated cardiac myocyte (**Supplemental Figure 1**). The full detection of a 500 second recording with 900 APs took approximately 1 second. By performing a linear fit of this runtime as a function of recordings duration, we found the SanPy algorithm takes approximately 0.01 s for each second of a 10 kHz recording. Timing was tested on a MacBook Pro (2021) with an Apple M1 Max CPU and 64 GB of memory. These timing results indicate that real-time use of SanPy for recordings well beyond 500 seconds should not be a hinderance. This also allows using the SanPy API on large datasets with hundreds of recordings to be rapidly batch-analyzed with the API.

### Analysis of recordings from neurons and cardiac myocytes

We have tested SanPy on several cell types including cardiac myocytes, dorsal root ganglion neurons, cortical layer V pyramidal neurons (provided by Niraj S. Desai, NINDS; Data not shown), and *in silico* stochastic cortical neurons. These tests were a critical step to ensure the provided detection parameter presets could be quickly fine-tuned for each of these cell types with potentially large heterogeneity in their membrane kinetics and the shape of APs. This was also important to ensure our detection algorithms yielded the correct analysis results.

In our previous work, an early version of SanPy was used to perform the analysis of current-clamp recordings of acutely isolated myocytes from the sinoatrial node of the heart (15). By examining anatomically distinct subpopulations of myocytes, we showed significant differences between superior and inferior sinoatrial node myocyte AP phenotypes including AP frequency and coefficient of variation, early diastolic duration (ms), and early diastolic depolarization rate (mV/s).

To further test SanPy on real-world data, here, we performed perforated patch current-clamp recordings in acutely isolated cardiac myocytes and used SanPy to analyze spontaneous APs. We compared within cell AP phenotypes in control versus in the present of the beta adrenergic receptor agonist ISO (**Figure 6a**). We found that ISO significantly increased AP frequency (Control 0.76 ± 0.72 Hz; ISO 1.17 ± 0.56 Hz, p = 0.0005) and half-width (Control 7.0 ± 0.5 ms; ISO 10.15 ± 0.83 ms, p < 0.0005) but had no significant effect on the early diastolic depolarization rate (Control 42.25 ± 45.4 mV/s; ISO 52.79 ± 38.26 mV/s, p = 0.12)(**Figure 6b**). These results indicate that the SanPy algorithms are robust and can detect the hypothesized changes in current-clamp analysis parameters.

**Figure 6.**
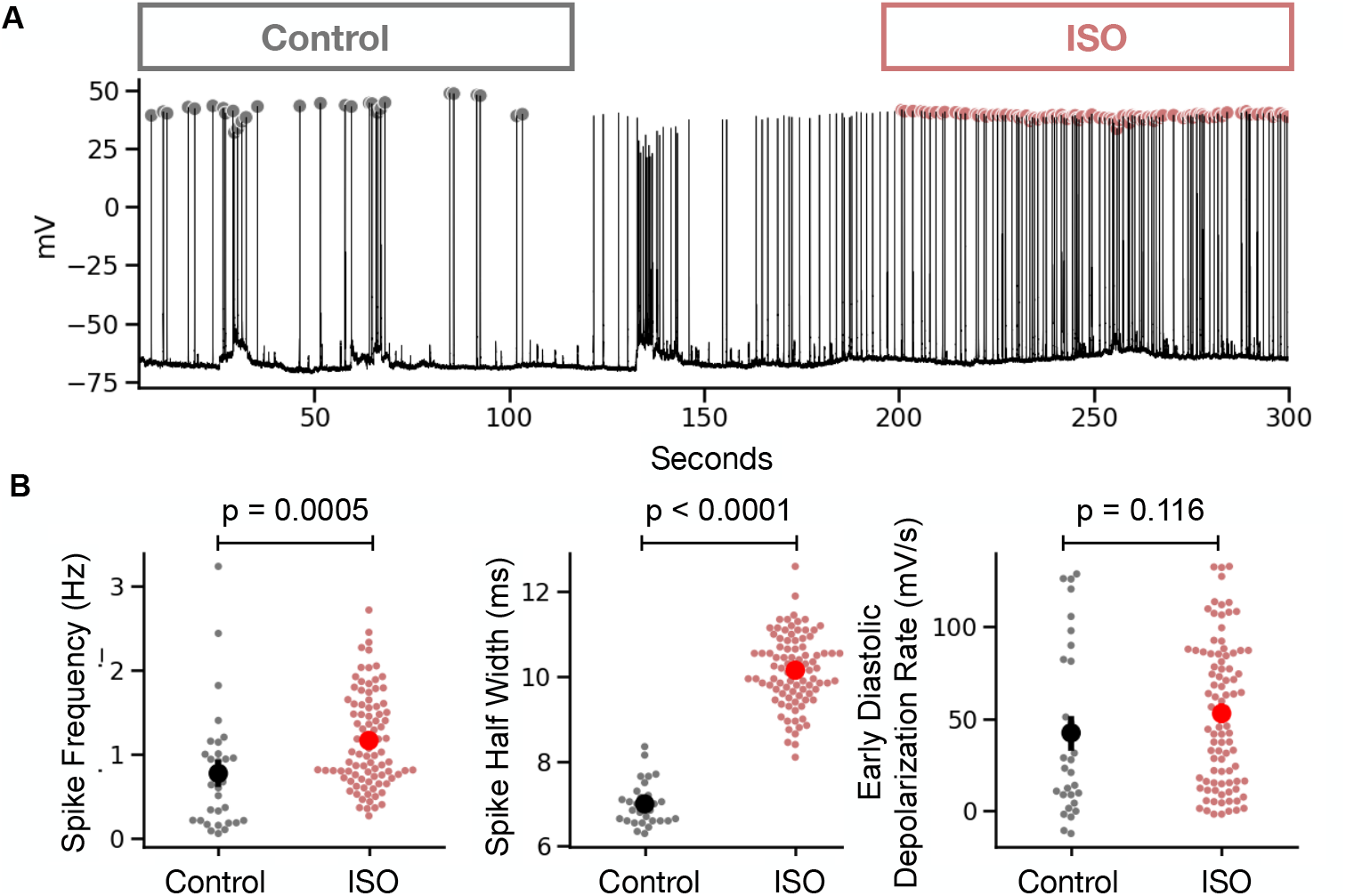
Example analysis of current-clamp recording from a cardiac myocyte. **(A)** Time-series plot of a perforated patch current-clamp recording in control and after bath application of ISO. Detected AP peaks are show as a circle with color encoding the condition (gray for control and red for ISO). **(B)** Summary statistics comparing analysis results in control versus ISO conditions. Statistical p values are for a Mann Whitney U test.

## Discussion

We present SanPy, a software analysis pipeline for whole-cell current-clamp recordings. SanPy can be used at multiple levels. Firstly and importantly, SanPy can be used as a desktop GUI application (Windows, macOS, and Linux) providing an intuitive and easy to use point-and-click interface that enables the analysis, exploration, and curation of electrophysiology data and analysis without requiring any complicated installation or programming. Secondly, the SanPy API allows all functionality to be programmatically scripted and extended, allowing programmers to fully extend SanPy. SanPy can be extended in several ways using the provided plugin architectures including custom file loaders, additions to the core analysis algorithms, and general-purpose interface plugins. Taken together, SanPy provides a solid foundation on which to perform a range of current-clamp electrophysiology analysis and with its extensibility will foster other research groups to build and share additional functionality.

Throughout its development, SanPy has been designed to fully satisfy the FAIR (Findability, Accessibility, Interoperability, and Reusability) methodology (32). SanPy is findable on GitHub and PyPi and is accessible with its easy-to-use desktop GUI. The modular design and API allow SanPy to interoperate with existing code bases and other analysis packages. Finally, SanPy ensures the reuse of data by clearly defining the meaning of both the detection parameters and analysis results.

SanPy has been tested on cardiac myocytes and several brain neuron types. There is nothing limiting the use of SanPy to any excitable cell type including, for example, skeletal and smooth muscle cells as well as endocrine cells such as insulin-releasing pancreatic β cells. To ensure SanPy can be used on any number of cell types, we have exposed all detection parameters to the end user. To reduce potential complexity, we provide pre-defined detection parameters for common cardiac and neuronal cell types that can be easily extended to other cell types. Finally, all detection parameters are documented and easily specified, saved, and loaded directly in the GUI.

To facilitate the curation of analysis results, SanPy flags APs when the detection parameters failed to find a signal. We believe this is a unique feature of SanPy, not found in other analysis software. For example, if the specified maximum half-with was shorter than the actual half-width in the recoded cell. A first key step in analysis curation is to check these detection errors to determine if the detection parameters need to be further adjusted. This can be done with the summary analysis plugin (**Table 1**). With this system of flagging errors, we are confident the end user will be able to identify the elusive and often lost false negatives in each analysis.

A key feature of the SanPy desktop GUI is the visualization of the analysis results in the context of the raw data. All plots are linked, selecting an analysis result such as AP time or peak amplitude will snap all other views to the raw data. This is critically important for several reasons including the curation of the analysis results by interrogating outliers. These outliers can be flagged to not be included in the results or the detection parameters can be fine-tuned to obtain more reliable results.

As a software ecosystem, SanPy provides a rich set of mechanisms by which it can be extended. This includes extensible file loaders, the addition of core analysis results, and a plugin architecture to create new and unique analysis and visualizations. With this, a group of researchers with differing skills can, for example, have programmers implement new analysis that can then be used by non-programmers in the point-and-click GUI.

Finally, we envision SanPy will be used in near-real-time during data acquisition to interrogate the quality of a recording and to perform customized analysis. By leveraging the plugin architecture, we envision that the analysis for experiments with unique requirements can be visualized and quantified during an actual experiment.

## Limitations and Future Directions

SanPy is optimized for the analysis of whole-cell current-clamp recording and the analysis of AP like events in excitable cells such as brain neurons and cardiac myocytes. It does not implement other common use cases such as the analysis of voltage-clamp recordings. This would include neuronal miniature excitatory and inhibitory post-synaptic currents or the probing of ion channel current activation and inactivation kinetics. There is nothing in the core functionality that limits the future implementation of this analysis, and we envision that the plugin architecture will facilitate the development of a range of these tools.

Although SanPy is currently limited in the number of raw data file types it can load, the provided file-loader functionality will allow this to be extended to any number of file formats. A key future direction is to incorporate both import and export of community-based file formats such as Neurodata Without Borders (NWB) and the Distributed Archives for Neurophysiology Data Integration (DANDI)(33).

Using our Python backend API, we have implemented a point-and-click desktop GUI. Because we have refined the backend to be easy to use, the same can be done to create web-based interfaces that run in a browser. In future versions of SanPy, this web-based GUI will greatly enhance the ability of groups of researchers to interact during data analysis and curation. This will also allow published results to be interactively viewed, further analysis to be performed, and models to be built by the greater research community.

## Acknowledgments

We are thankful for feedback from members of the Santana lab and the UC Davis community. We thank Niraj S. Desai (NINDS) for providing sample Matlab file-format electrophysiology recordings of cortical Layer-V pyramidal neurons. We are indebted to open-source developers from around the world including the thousands that have contributed to Python packages. We are particularly thankful for the work done by Scott W. Harden on pyAbf. This work was supported by grants from the US National Institutes of Health to R.H.C. (1RF1MH123206) and to L.F.S. (1R01HL144071 and 1OT2OD026580).

