## Supplemental Material for "SanPy: A whole-cell electrophysiology analysis pipeline"

All tables and recipes included here are available online in the SanPy documentation.

**Supplemental Table 1.** Definition of detection parameters.
See: https://cudmore.github.io/SanPy/methods/#detection-parameters

**Supplemental Table 2.** Analysis results for each detected AP.
See: https://cudmore.github.io/SanPy/methods/#analysis-output-full

**Supplemental Recipe 1.**
See: https://cudmore.github.io/SanPy/api/writing-a-file-loader/

**Supplemental Recipe 2.**
See: https://cudmore.github.io/SanPy/api/writing-new-analysis/

**Supplemental Recipe 3.**
See: https://cudmore.github.io/SanPy/api/writing-a-plugin/

**Supplemental Figure 1.** Analysis of runtime for AP detection

**Supplemental Table 1.** Definition of detection parameters.
See: https://cudmore.github.io/SanPy/methods/#detection-parameters

| Parameter | Default Value | Units | Human Readable | Description |
|---|---|---|---|---|
| detectionName | default | | Detection Preset Name | The name of detection preset |
| detectionType | dvdt | | Detection Type | Detect using derivative (dvdt) or membrane potential (mV) |
| dvdtThreshold | 20 | dVdt | dV/dt Threshold | dV/dt threshold for a spike, will be backed up to dvdt_percentOfMax and have xxx error when this fails |
| mvThreshold | -20 | mV | mV Threshold | mV threshold for spike AND minimum spike mV when detecting with dV/dt |
| startSeconds | | s | Start(s) | Start seconds of analysis |
| stopSeconds | | s | Stop(s) | Stop seconds of analysis |
| condition | “” | String | Condition | Condition |
| dvdt_percentOfMax | 0.1 | Percent | dV/dt Percent of max | For dV/dt detection, the final TOP is when dV/dt drops to this percent from dV/dt AP peak |
| onlyPeaksAbove_mV | | mV | Accept Peaks Above (mV) | Only accept APs with peaks above this value (mV) |
| onlyPeaksBelow_mV | | mV | Accept Peaks Below (mV) | Only accept APs below this value (mV) |
| doBackupSpikeVm | False | Boolean | Backup Vm Spikes | If true, APs detected with just mV will be backed up until Vm falls to xxx |
| refractory_ms | 170 | ms | Minimum AP interval (ms) | APs with interval (wrt previous AP) less than this will be removed |
| peakWindow_ms | 100 | ms | Peak Window (ms) | Window after TOP (ms) to search for AP peak (mV) |
| dvdtPreWindow_ms | 10 | ms | dV/dt Pre Window (ms) | Window (ms) to search before each TOP for real threshold crossing |
| dvdtPostWindow_ms | 20 | ms | dV/dt Post Window (ms) | Window (ms) to search after each AP peak for minimum in dv/dt |
| mdp_ms | 250 | ms | Pre AP MDP window (ms) | Window (ms) before an AP to look for MDP |
| avgWindow_ms | 5 | ms | MDP averaging window (ms) | Window (ms) to calculate MDP (mV) as a mean rather than mV at single point for MDP |
| lowEddRate_warning | 8 | EDD slope | EDD slope warning | Generate warning when EED slope is lower than this value. |
| halfHeights | [10, 20, 50, 80, 90] | | AP Durations (%) | AP Durations as percent of AP height (AP Peak (mV) - TOP (mV)) |
| halfWidthWindow_ms | 200 | ms | Half Width Window (ms) | Window (ms) after TOP to look for AP Durations |
| medianFilter | 0 | points | Median Filter Points | Number of points in median filter, must be odd, 0 for no filter |
| SavitzkyGolay_pnts | 5 | points | SavitzkyGolay Points | Number of points in Savitzky-Golay filter, must be odd, 0 for no filter |
| SavitzkyGolay_poly | 2 | | SavitzkyGolay Poly Deg | The degree of the polynomial for Savitzky-Golay filter |
| verbose | FALSE | Boolean | Verbose | Verbose Detection Reporting |

**Supplemental Table 2.** Analysis results for each detected AP.
See: https://cudmore.github.io/SanPy/methods/#analysis-output-full

| Name | type | Description |
|---|---|---|
| analysisDate | str | Date of analysis in yyyymmdd ISO 8601 format. |
| analysisTime | str | Time of analysis in hh:mm:ss 24 hours format. |
| modDate | str | Modification date if AP is modified after detection. |
| modTime | str | Modification time if AP is modified after detection. |
| analysisVersion | str | Analysis version when analysis was run. See sanpy.analysisVersion |
| interfaceVersion | str | Interface version string when analysis was run. See sanpy.interfaceVersion |
| file | str | Name of raw data file analyzed |
| detectionType | | Type of detection, either vm or dvdt. See enum sanpy.bDetection.detectionTypes |
| cellType | str | User specified cell type |
| sex | str | User specified sex |
| condition | str | User specified condition |
| sweep | int | Sweep number of analyzed sweep. Zero based. |
| epoch | int | Stimulus epoch number the spike occurred in. Zero based. |
| epochLevel | float | Epoch level (DAC) stimulus during the AP threshold. |
| sweepSpikeNumber | int | AP number within the sweep. Zero based. |
| spikeNumber | int | AP number across all sweeps. Zero based. |
| include | bool | Boolean indication include or not. Can be set by user/programmatically after analysis. |
| userType | int | Integer indication user type. Can be set by user/programmatically after analysis. |
| errors | list | List of dictionary to hold detection errors for this AP |
| dvdtThreshold | float | AP Threshold in derivative dv/dt |
| mvThreshold | float | AP Threshold in primary recording mV |
| medianFilter | int | Median filter to generate filtered vm and dvdt. Value 0 indicates no filter. |
| halfHeights | list | List of int to specify half-heights like [10, 20, 50, 80, 90]. |
| thresholdPnt | int | AP threshold point |
| thresholdSec | float | AP threshold seconds |
| thresholdVal | float | Value of Vm at AP threshold point. |
| thresholdVal_dvdt | float | Value of dvdt at AP threshold point. |
| dacCommand | float | Value of DAC command at AP threshold point. |
| peakPnt | int | AP peak point. |
| peakSec | float | AP peak seconds. |
| peakVal | float | Value of Vm at AP peak point. |
| peakHeight | float | Difference between peakVal minus thresholdVal. |
| timeToPeak_ms | float | Time to peak (ms) after TOP. |
| preMinPnt | int | Minimum before an AP taken from predefined window. |
| preMinVal | float | Minimum before an AP taken from predefined window. |
| preLinearFitPnt0 | int | Point where pre linear fit starts. Used for EDD Rate |

| | | |
|---|---|---|
| preLinearFitPnt1 | int | Point where pre linear fit stops. Used for EDD Rate |
| earlyDiastolicDuration_ms | float | Time (ms) between start/stop of EDD. |
| preLinearFitVal0 | float | |
| preLinearFitVal1 | float | |
| earlyDiastolicDurationRate | float | Early diastolic duration rate, the slope of the linear fit between start/stop of EDD. |
| lateDiastolicDuration | float | Depreciated |
| preSpike_dvdt_max_pnt | int | Point corresponding to peak in dv/dt before an AP. |
| preSpike_dvdt_max_val | float | Value of Vm at peak of dv/dt before an AP. |
| preSpike_dvdt_max_val2 | float | Value of dv/dt at peak of dv/dt before an AP. |
| postSpike_dvdt_min_pnt | int | Point corresponding to min in dv/dt after an AP. |
| postSpike_dvdt_min_val | float | Value of Vm at minimum of dv/dt after an AP. |
| postSpike_dvdt_min_val2 | float | Value of dv/dt at minimum of dv/dt after an AP. |
| isi_pnts | int | Inter-Spike-Interval (points) with respect to previous AP. |
| isi_ms | float | Inter-Spike-Interval (ms) with respect to previous AP. |
| spikeFreq_hz | float | AP frequency with respect to previous AP. |
| cycleLength_pnts | int | Points between APs with respect to previous AP. |
| cycleLength_ms | int | Time (ms) between APs with respect to previous AP. |
| diastolicDuration_ms | float | Time (ms) between minimum before AP (preMinPnt) and AP time (thresholdPnt). |
| widths_10 | int | Width (ms) at half-height 10 %. |
| widths_20 | int | Width (ms) at half-height 20 %. |
| widths_50 | int | Width (ms) at half-height 50 %. |
| widths_80 | int | Width (ms) at half-height 80 %. |
| widths_90 | int | Width (ms) at half-height 90 %. |

**Supplemental Recipe 1.**
See: https://cudmore.github.io/SanPy/api/writing-a-file-loader/

### How to write a custom SanPy file loader.

1) Derive a new class from sanpy.fileloaders.fileLoader_base.

2) Specify the file extension you want to load with `loadFileType = 'your_file_extension'`

3) In a `loadFile()` member function, load your raw data file

4) Call `self.setLoadedData(...)` with the results.

5) Place your file loader py file in the `<User>/Documents/SanPy/file loaders` folder.

6) Run SanPy and make sure it works!

**Coming Soon.** We will provide unit testing for user file loaders.

Here is some sample code to get started, this is taken from the SanPy CSV file loader fileLoader_csv.

```
import sanpy.fileloaders.fileLoader_base as fileLoader_base

class fileLoader_csv(fileLoader_base):
    loadFileType = 'csv'

    def loadFile(self):
        """Load file and call setLoadedData().

        Use self.filepath for the file path to load
        """
```

The function signature for `setLoadedData` is as follows. There are only two required parameters and a number of optional parameters.

```
    def setLoadedData(self,
        sweepX : np.ndarray,
        sweepY : np.ndarray,
        sweepC : Optional[np.ndarray] = None,
        recordingMode : recordingModes = recordingModes.iclamp,
        xLabel : str = '',
        yLabel : str = ''):

        """
        Parameters
        ----------
        sweepX : np.ndarray
            Time values
        sweepY : np.ndarray
            Recording values, mV or pA
        sweepC : np.ndarray
            (optional) DAC stimulus, pA or mV
        recordingMode : recordingModes
            (optional) Defaults to recordingModes.iclamp)
        xLabel : str
            (optional) str for x-axis label
        yLabel : str
            (optional) str for y-axis label
        """
```

**Supplemental Recipe 2.**
See: https://cudmore.github.io/SanPy/api/writing-new-analysis/

### How to extend the analysis of SanPy with user specified analysis

The core analysis algorithm of SanPy can be easily extended by the user.

- Derive a class from sanpy.user_analysis.baseUserAnalysis
- In the member function `defineUserStats()`, define the name of your new analysis with addUserStat()
- In the member function `run()`, for each spike in the analysis, calculate your new stat and set the value with `setSpikeValue()`.

Here is a simple example to get you started.

```
class exampleUserAnalysis(baseUserAnalysis):
    """
    An example user defined analysis.

    We will add 'User Time To Peak (ms)', defines as:
        For each AP, the time interval between spike threshold and peak

    We need to define the behavior of two inherited functions

    1) defineUserStats()
        add any numner of user stats with
            addUserStat(human_name, internal_name)

    2) run()
        Run the analysis you want to compute.
        Add the value for each spike with
            setSpikeValue(spike_index, internal_name, new_value)
    """

    def defineUserStats(self):
        """Add your user stats here."""
        self.addUserStat("User Time To Peak (ms)", "user_timeToPeak_ms")

    def run(self):
        """This is the user code to create and then fill in
            a new name/value for each spike."""

        # get filtered vm for the entire trace
        # filteredVm = self.getFilteredVm()

        for spikeIdx, spikeDict in enumerate(self.ba.spikeDict):

            # add time to peak
            thresholdSec = spikeDict["thresholdSec"]
            peakSecond = spikeDict["peakSec"]
            timeToPeak_ms = (peakSecond - thresholdSec) * 1000

            # assign to underlying bAnalysis
            self.setSpikeValue(spikeIdx, "user_timeToPeak_ms", timeToPeak_ms)
```

**Supplemental Recipe 3.**
See: https://cudmore.github.io/SanPy/api/writing-a-plugin/

### How to write a SanPy plugin.

1) Derive a class from sanpy.interface.plugins.sanpyPlugin

2) Give you plugin a name by defining the static property `myHumanName = 'Nice name for your plugin`.

3) Build your user interface in a `plot()` member function.

4) Have your plugin respond to the main interface by reploting in a `replot()` member function. This is to enable your plugin to respond to different pre-defined interface changes, see below.

5) Place you new plugin py file in the `<user>Documents/SanPy/plugins` folder

6) Run SanPy and it will append you plugin to the list of available plugins in the `Plugins Menu`.

**Coming soon.** We will provide unit tests to ensure new plugins is working.

Here is a template to get started. This is the same as gets installed in the User plugin folder file `exampleUserPlugin.py`.

```
from sanpy.interface.plugins import sanpyPlugin

class exampleUserPlugin1(sanpyPlugin):
    """
    Plot x/y statistics as a scatter

    Get stat names and variables from sanpy.bAnalysisUtil.getStatList()
    """
    myHumanName = 'Example User Plugin 1'

    def plot(self):
        """Create the plot in the widget (called once).
        """

        # embed a matplotlib axis (self.axs)
        self.mplWindow2() # assigns (self.fig, self.axs)

        # plot a white line with raw data
        self._lineRaw, = self.axs.plot([], [], '-w', linewidth=0.5)

        # plot red circles with spike threshold
```

```
        self._lineDetection, = self.axs.plot([], [], 'ro')

    def replot(self):
        """Replot the widget. Usually when the file is switched
        """
        # get the x/y values from the recording
        sweepX = self.getSweep('x')
        sweepY = self.getSweep('y')

        # update plot of raw data
        self._lineRaw.set_data(sweepX, sweepY)

        # update plot of spike threshold
        thresholdSec = self.ba.getStat('thresholdSec')
        thresholdVal = self.ba.getStat('thresholdVal')
        self._lineDetection.set_data(thresholdSec, thresholdVal)

        # make sure the matplotlib axis auto scale
        self.axs.relim()
        self.axs.autoscale_view(True,True,True)

        # plt.draw()
        self.static_canvas.draw()
```

**Supplemental Figure 1.** Analysis of runtime for AP detection

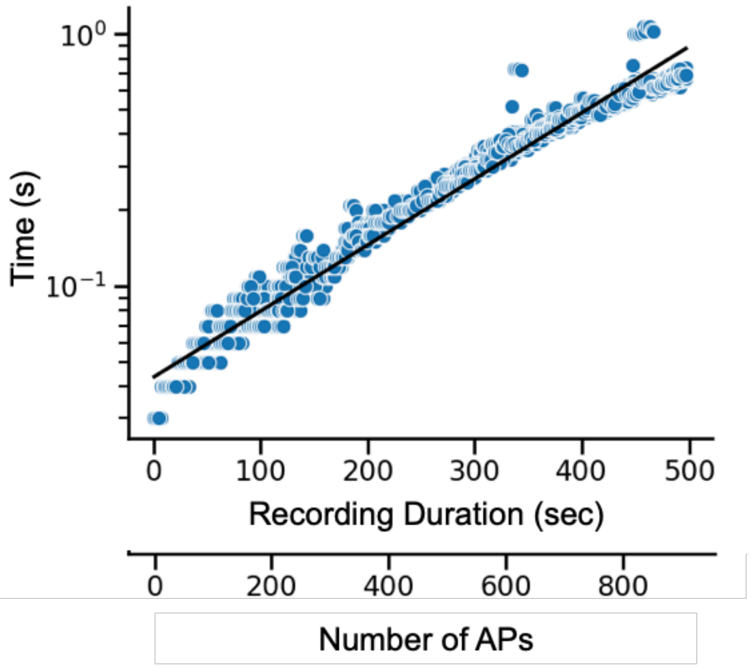


**Supplemental Figure 1. Analysis of runtime for AP detection**. Plot of the time to execute AP detection versus the recording duration analyzed and corresponding number of APs detected. Each symbol (circle) represents one run of the detection algorithm for a given recording duration and number of APs. Y-axis is a log scale. Exponential fit is show as a line (black).
